# The first recorded incidence of *Deinococcus radiodurans* R1 biofilm formation and its implications in heavy metals bioremediation

**DOI:** 10.1101/234781

**Authors:** Sudhir K. Shukla, T. Subba Rao

## Abstract

Radiation tolerant *Deinococcus radiodurans* R1 is reported to be a potential bacterium for the treatment of low level active wastes. So far there are no reports on the biofilm producing capability of *D. radiodurans* and heavy metal biosorption. In this study, it was observed that a recombinant *D. radiodurans* strain with a plasmid harbouring *gfp* and *kan^R^* has formed significant biofilm (~10 μm thick). Analysis of biofilm matrix components produced by *D. radiodurans* showed that the matrix consisted primarily of proteins and carbohydrates with a little amount of extracellular DNA (eDNA). Further, studies showed that *D. radiodurans* biofilm formation was enhanced at higher concentrations (up to 25 mM) of Ca^2+^. Further studies on *D. radiodurans* biofilm showed that Ca^2+^ enhanced significant biosorption of the heavy metals (Co, Ni). In the presence of 25 mM Ca^2+^, the *D. radiodurans* (Kan^r^) biofilm showed 35% and 25% removal of Co^2+^ and Ni^2+^ respectively. While in the absence of Ca^2+^, *D. radiodurans* biofilm showed relatively low biosorption of Co (7%) and Ni (3%). Ca^2+^ also significantly enhanced exopolysaccharide (EPS) production in the biofilm matrix. This infers that EPS could have mediated the heavy metal biosorption. This study signifies the potential use of *D. radiodurans* biofilm in the remediation of radioactive waste components.

**Significance and Impact of this Study:** This is the first ever recorded study on the *Deinococcus radiodurans* R1 biofilm. This organism, being the most radioresistant micro-organism ever known, has always been speculated as a potential bacterium to develop a bioremediation process for radioactive heavy metal contaminants. However, the lack of biofilm forming capability proved to be a bottleneck in developing such technology. This study records the first incidence of biofilm formation in a recombinant *D. radiodurans*, serendipitously, and also discusses its implications in removal of heavy-metals, such as Co and Ni.

## Introduction

Biofilms are a preferred mode of life by most bacteria in nature. The biofilm is made up of a thin layer of microbes where matrices of self-made biopolymer molecules encase the cells. Biofilm mode of living provides a bacterial cell protection from environmental, chemical and physical stresses (Costerton *et al.* 1987); (Donlan 2002; Das *et al.* 2012). Biofilms apart from enhancing the organisms survival, also aids in improving the rate and the extent of contaminant transformation as compared to that of pure and planktonic cultures (Mangwani *et al.* 2014a). Remediation using microorganism (termed as bioremediation) is an emerging *in situ* technology for the clean-up of environmental pollutants. The economical factor and inefficiency of some physicochemical remediation methods has made this biological treatment method as an improved alternative (Paul *et al.* 2005). Before employing any biofilm forming bacterium for developing bioremediation process, a detailed study needs to be carried out. Previous studies have shown that biofilm parameters and EPS composition greatly affects the pollutant degradation and remediation potential (Mangwani *et al.* 2014b; Mangwani *et al.* 2016). Apart from intrinsic factors such as biofilm formation and quorum sensing (Mangwani *et al.* 2012), extrinsic factors such as the presence of Ca^2+^ also modulate the biofilms (Mangwani *et al.* 2014b; Shukla and Rao 2014) and in turn augment bioremediation capability.

The bioremediation potential of *D. radiodurans* has gained importance in recent times. *D. radiodurans* has an extraordinary capability of surviving under high radiation stress (Battista 1997), even in low nutrient conditions (Shukla *et al.* 2014b). Engineered *D. radiodurans* cells have been used to detoxify mercury, degrade toluene (Brim *et al.* 2000) and reduce chromium (Brim *et al.* 2006). However, use of planktonic cells for bioremediation of heavy metals makes the downstream process costly or less efficient. Biofilm-mediated bioremediation processes are more efficient as compared to the processes mediated by their planktonic counterparts. To use any bacterium for bioremediation purpose, the knowledge about its biofilm production characteristics is a prerequisite. Till date, there is not a single report on biofilm formation of *D. radiodurans*. A genetically engineered strain of *D. radiodurans*, expressing *gfp* plasmid (GenBank accession no. KF975402), was found to produce biofilm in our lab. However, the exact mechanism behind the acquired biofilm forming capability is not yet clear and is under investigation. In this study, we focus only on the characterisation of the *D. radiodurans* biofilm and its implications in the bioremediation of heavy metals.

Over the past few decades, rapid growth of chemical industries has enhanced the heavy metal release into the environment, leading to contamination of air, water and atmosphere (Akpor and Muchie 2010). These heavy metals need to be converted from a toxic form to a lesser hazardous form or its bioavailability should be decreased (Wall and Krumholz 2006). Naturally occurring and comparatively less abundant cobalt and nickel metals have drawn the attention due to surge in their anthropogenic activities and the potential hazards due to any accidental release into the environment (Shukla *et al.* 2017). The intake of these ions can cause detrimental health hazards (Keith *et al.* 2013). The chemical methods used for radioactive heavy metal precipitation has its limitations due to high cost and low feasibility (Shukla *et al.* 2014a). Apart from the higher cost burden, most of such chemical methods for the removal of radioactive heavy metals end up with the generation of a large amounts of sludge and also incomplete removal of metal ions (Gadd 2010).

To overcome these disadvantages, bioremediation approaches using heavy metal resistant microorganisms like bacteria and fungus are promising due to their low cost as well as feasibility for *in situ* applications (Yan and Viraraghavan 2003; Vijayaraghavan *et al.* 2005; Pal *et al.* 2006). Bioremediation using microbes can be a very promising and more efficient approach as microbes are nature’s creative recyclers (N’Guessan *et al.* 2008). It was inferred that the bacteria, fungus and algae isolated from heavy metal polluted areas are the ideal candidates for the bioremediation of heavy metals (Colin *et al.* 2012). A number of studies have been carried out in order to utilize microorganisms as metal bio-sorbents (Vijayaraghavan *et al.* 2005; Pal *et al.* 2006; Paraneeiswaran *et al.* 2014), however, reports on biofilm mediated-biosorption of heavy metals are scarce.

In this study, *D. radiodurans* biofilm was characterised by using classical crystal violet assay and confocal laser scanning microscopy. We also studied the effect of Ca^2+^ on biofilm formation and its EPS composition. Apart from biofilm characterisation, it was also investigated whether this recombinant strain of *D. radiodurans* can be used for biofilm mediated remediation of heavy metals.

## Materials & Methods

### Microorganism and culture conditions

*Deinococcus radiodurans* R1 wild type (DR1-WT) and *gfp*-harbouring, biofilm forming strain of *D. radiodurans* (DR1-Kan^r^) was used in this study. Both the strain of *D. radiodurans* were maintained on TGY (tryptone-glucose-yeast extract) medium consisting of 5 g tryptone, 3 g yeast extract, 1 g glucose and 1.5% agar was added to prepare solid medium for sub-culturing and culture purity study. The inoculated TGY broth cultures were incubated at 30°C in an orbital shaker at 100 rpm until mid-log phase was reached. Kanamycin antibiotic was added in the TGY at final concentration of 5 μg/mL for the growth of *D. radiodurans* (Kan^r^).

### Quantitative biofilm assay

Biofilm assay was performed to assess biofilm production by *D. radiodurans* strains. Biofilm quantification was done by an improved method of classical crystal violet assay (Shukla and Rao 2017). The overnight grown cultures of the *D. radiodurans* strains in TGY were diluted 1:40 in sterile TGY medium and added to the pre-sterilized flat bottom polystyrene 96 well micro-titre plates. After the prescribed time as per the experimental plan, biofilms were gently washed with 200 μl of sterile phosphate buffer saline (PBS) and stained with 0.2% crystal violet solution for 5 min. Thereafter, the excess crystal violet was decanted and biofilms were washed again with PBS twice. Crystal violet was solubilised with 95% alcohol and biofilm quantification was done in terms of absorbance at 570 nm using a multimode micro-plate reader (BioTek, USA).

To study the effect of calcium on *D. radiodurans* (Kan^r^) biofilm growth, the biofilm was grown as described above but in the presence of different concentrations of CaCl_2_ in the range of 1–50 mM in TGY broth for 48 h. *D. radiodurans* (Kan^r^) in TGY broth without Ca^2+^ served as control, and TGY broth alone considered as blank. Biofilm growth was monitored using crystal violet assay as described above.

### Confocal laser scanning microscopy (CSLM) studies and image analysis

Biofilms grown on pre-sterilized microscopic glass slides were studied using CSLM (Mangwani *et al.* 2014b). The overnight grown culture of the *D. radiodurans* (Kan^r^) was diluted 1:40 in sterile TGY medium and added to the pre-sterilized flat bottom 6 well plates. Sterile glass slides were immersed into the growth medium as a substratum for biofilm growth. The petri-plates were incubated at 80 rpm for 48 h at 30°C. The glass slides were gently washed with PBS to remove loosely attached cells and stained with 0.2 % acridine orange for 5 min, thereafter the slide is thoroughly washed with PBS. A thin cover slide was mounted over the stained biofilm and observed by keeping the slide upside down on objective lens of the confocal laser scanning microscope (TCS SP2 AOBS) equipped with DM IRE 2-inverted microscope (Leica Microsystems, Germany). Image stacks were collected from 20 random points of the biofilms in order to get an accurate mean value of the biofilm parameters. Quantification of the biofilm parameters (average thickness, maximum thickness, total biomass, roughness coefficient, and surface to biovolume ratio) was done by COMSTAT program written as a script in MATLAB 5.2 software (Heydorn *et al.* 2000). Each experiment was repeated three times to have statistically significant data.

### EPS extraction and quantification of biofilm matrix components

*Deinococcus radiodurans* (Kan^r^) biofilm was grown for 48 h on glass slides immersed in 20 ml of TGY. After 48 h, planktonic cells were aspirated and biofilm was gently washed twice with PBS, and then remaining biofilm was scrapped and collected in 5 ml of PBS. Biofilm was disintegrated by gentle vortexing using glass beads. A 5 ml of the biofilm sample was centrifuged at 8000 rpm and 4°C for 30 min. Supernatant was collected and mixed with double volume of 90% chilled ethanol and kept at 4°C overnight. EPS was collected by centrifugation at 10000 rpm and 4°C for 10 min. The supernatant was discarded and pellet was collected and dried at 60°C to remove ethanol. Pellet was resuspended in 100 μl PBS buffer. The protein and eDNA content in the resuspension was quantified using Qubit Fluorimeter (Invitrogen, Carlsbad, CA, USA), the quantification protocol was followed as described by the vendor (Beaudet *et al.* 2007; Bajrami *et al.* 2009). Glucose concentration as a measure of polysaccharide content was quantified by the Dubois method as described elsewhere (Paraneeiswaran *et al.* 2015).

### Biofilm-mediated heavy metal removal

*Deinococcus radiodurans* (Kan^r^) was grown in TGY broth in the presence of Ca^2+^ (25 mM Ca^2+^) and incubated for 48 h. Biofilm with 25 mM Ca^2+^ was taken in a total volume of 200 μl sterile TGY broth in 96 well microtitre plate. After incubation, the heavy metals such as Cobalt and Nickel were added. The TGY broth with heavy metals (Co^2+^ and Ni^2+^) was considered as control. After 24 h of incubation, the supernatants were collected from the well of plates. The supernatant was filtered through 0.22 μM membrane filter. Then the concentrations of heavy metals (Co^2+^ and Ni^2+^) present in the collected supernatants was analysed using Inductively Coupled Plasma-Atomic Emission Spectroscopy (ICP-AES, model: Ultima-2, Horiba JovinYvon, France).

### Statistical analysis

All data are expressed as mean standard deviation (SD) of the triplicate experimental data. A two-tailed Student’s t-test was used to determine the differences in biofilm formation between the control and each group. The P value of < 0.05 was taken as significant.

## Results

### Biofilm formation of by *D. radiodurans*

Biofilm growth capabilities of *D. radiodurans* R1 wild type and *D. radiodurans* (Kan^r^) were studied in two growth media, *i.e.*, LB and TGY by using classical crystal violet assay in 96 well microtitre plates (Fig. 1). Biofilm production was quantified at different time intervals at 24, 48, and 72 h. Results showed that *D. radiodurans* (Kan^r^) formed a good amount of biofilm as compared to *D. radiodurans* R1 wild type in both the growth medium. As shown in Fig. 1, a gradual increase was observed in *D. radiodurans* (Kan^r^) biofilm formation up to 72 h when grown in LB medium. On the other hand, maximum biofilm growth was observed after 48 h when the biofilm was grown in the presence of TGY medium. However, there was no significant difference in biofilm production at 48 h and 72 h. It can also be observed in Fig. 1A and 1B, that *D. radiodurans* (Kan^r^) biofilm growth was much better in TGY medium when compared to LB at all the time intervals monitored *i.e.*, 24, 48 and 72 h. Biofilm growth in TGY after 24, 48 and 72 h was found to be 44.8%, 75.9% and 34.34% higher as compared to LB medium. This observation indicated that TGY is the medium of choice for all biofilm related experiments.

**Figure 1.**
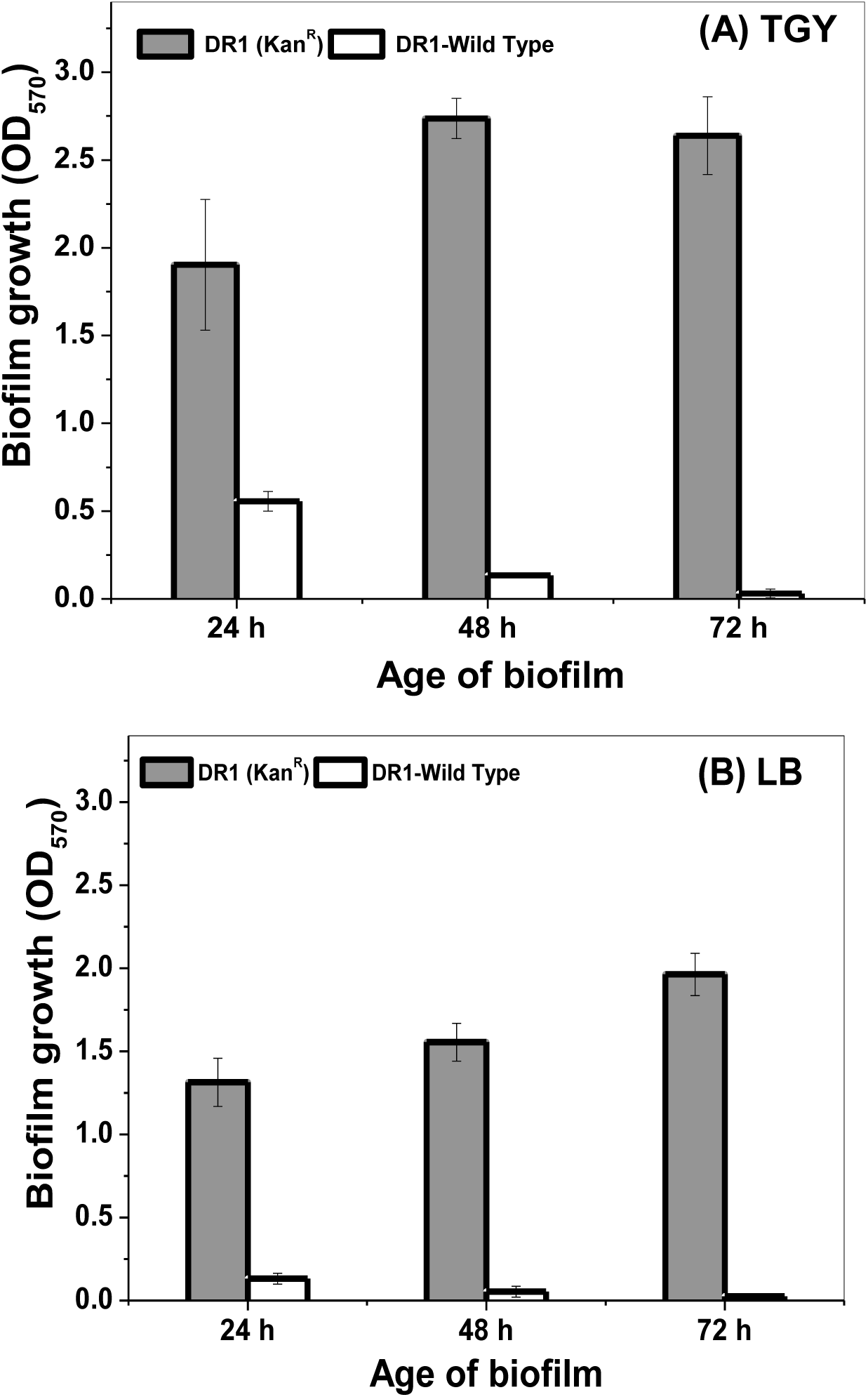
Biofilm formation by *D. radiodurans* R1 wild type and *D. radiodurans* (Kan^r^) bacterium in two different growth medium; (A) Tryptone Glucose Yeast extract (TGY), and (B) Luria Bertani (LB). Biofilms were evaluated after 24, 48 and 72 h of growth. Error bars are shown as ± SD.

**Figure 2:**
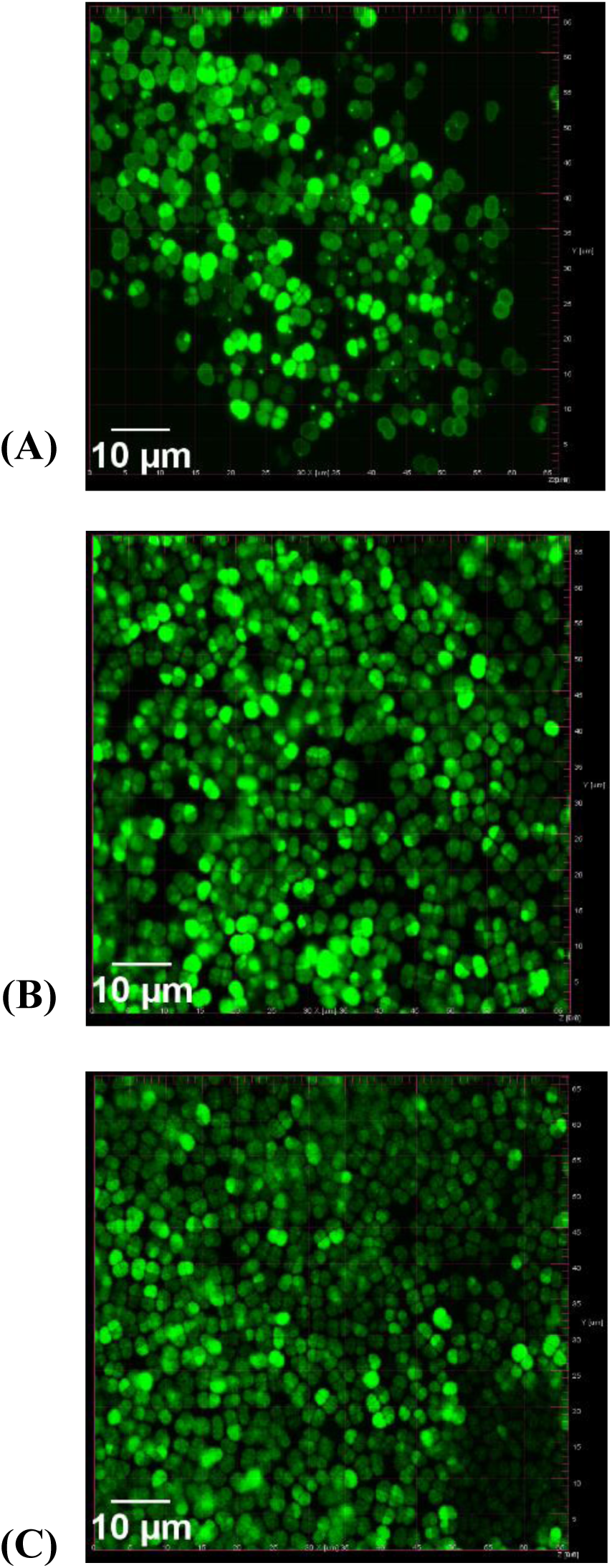
Confocal Laser Scanning Microscopic images of different days old *D. radiodurans* biofilm. (A) 24 h old biofilm (B) 48 h old biofilm (C) 72 h old biofilm (Kan^r^). (scale bar 10 μm).

### Confocal laser scanning microscopic study of *D. radiodurans* (Kan^r^) biofilm

Confocal laser scanning microscopic study of the *gfp* tagged *D. radiodurans* (Kan^r^) biofilm was done at 24 h, 48 h and 72 h to characterize the biofilm parameters. As shown in Table 1, biofilm parameters of *D. radiodurans* biofilm significantly changed over the time. Maximum biomass, average thickness of biofilm and average colony size at substratum were observed at 48 h. Total biomass μm^3^/μm^2^) at 24 h, 48 h and 72 h was found to be 0.53 ± 0.34, 1.63 ±1.08 and 0.87 ±0.65 respectively. Total biomass is positively correlated with the average thickness of biofilms. For *D. radiodurans* (Kan^r^) biofilm, *R*\* was found to be lowest at 48 h and surface to bio-volume ratio was found to be maximum at 48 h as compared to 24 and 72 h.

**Table 1:** Structural parameters of *D. radiodurans* (Kan^r^) biofilms in TGY medium at different days. Biofilm parameters were quantified from the CSLM images stacks.

| Biofilm parameters | Age of *D. radiodurans* (Kan<sup>r</sup>) biofilm |  |  |
| --- | --- | --- | --- |
|  | 24 h | 48 h | 72 h |
| Total biomass ($\mu\text{m}^3/\mu\text{m}^2$) | $0.53 \pm 0.39$ | $1.63 \pm 1.08$ | $0.87 \pm 0.65$ |
| Maximum thickness ($\mu\text{m}$) | $8.34 \pm 2.00$ | $9.12 \pm 0.50$ | $10.99 \pm 2.42$ |
| Avg. Thickness | $2.22 \pm 2.33$ | $4.03 \pm 2.54$ | $3.19 \pm 2.85$ |
| Avg. Colony size at substratum ($\mu\text{m}^2$) | $3.39 \pm 1.40$ | $4.14 \pm 2.47$ | $3.71 \pm 1.76$ |
| Avg. Colony volume at substratum ($\mu\text{m}^3$) | $1335.16 \pm 1119.19$ | $5267.83 \pm 5327.12$ | $2387.15 \pm 2467.58$ |
| Surface to volume ratio ($\mu\text{m}^2/\mu\text{m}^3$) | $3.24 \pm 0.22$ | $4.72 \pm 0.93$ | $4.31 \pm 1.3$ |
| Roughness coefficient | $1.55 \pm 0.32$ | $1.00 \pm 0.59$ | $1.34 \pm 0.47$ |

### Effect of Ca^2+^ on *D. radiodurans* biofilm formation and EPS production

The effect of Ca^2+^ on *D. radiodurans* (Kan^r^) biofilm growth was studied. The result showed that there was a significant increase in the *D. radiodurans* (Kan^r^) biofilm production with increasing concentration of Ca^2+^, when tested in the range of 1 - 50 mM (Fig. 3). Calcium enhanced the biofilm growth in a dose dependent manner. Optimum Ca^2+^ concentration for enhancing biofilm production was found to be 25 mM without causing substantial turbidity in TGY medium.

**Figure 3:**
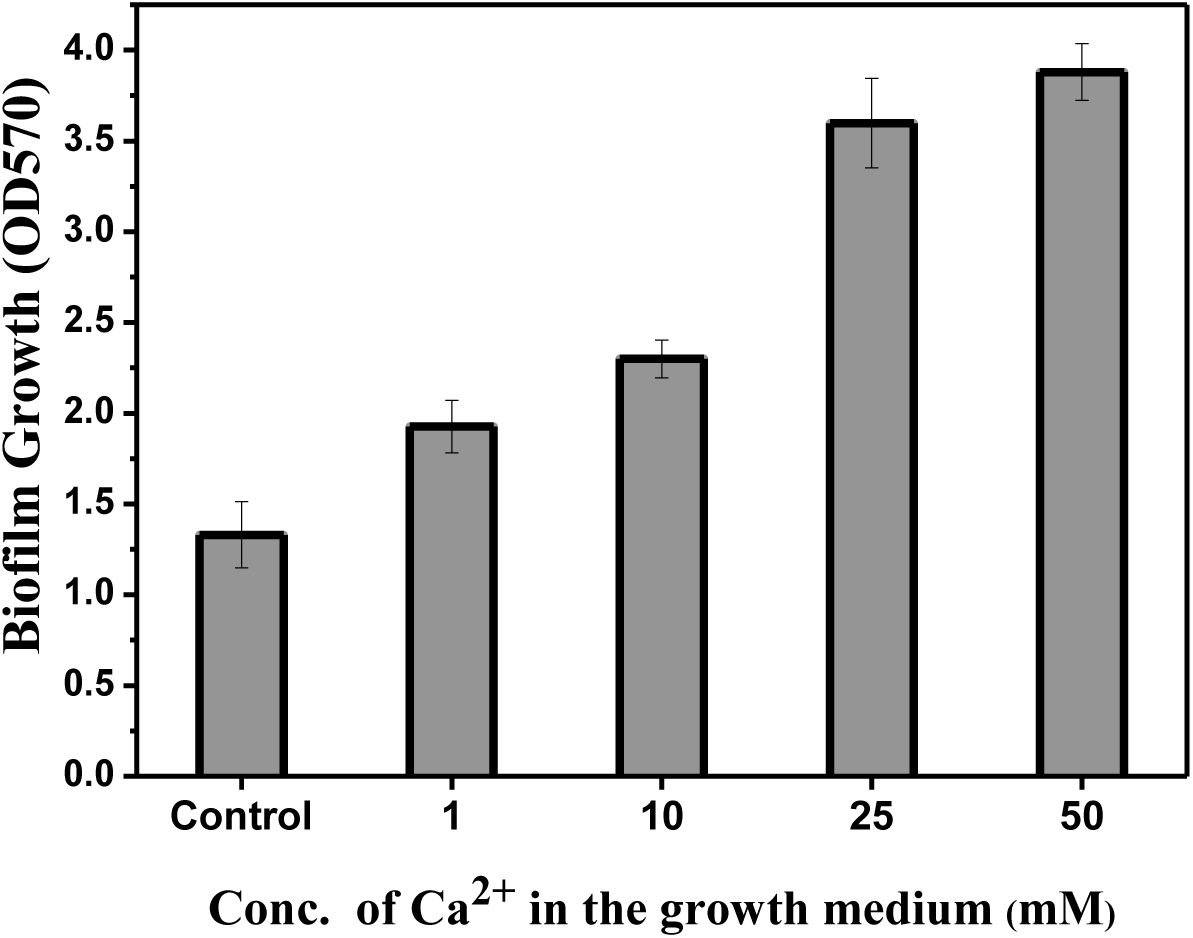
Effect of different concentrations of calcium on *D. radiodurans* (Kan^r^) biofilm growth. Biofilms were evaluated after 48 h of growth. Error bars are shown as ± SD.

To assess the effect of Ca^2+^ on EPS matrix the *D. radiodurans* (Kan^r^) biofilm was grown with and without Ca^2+^ and its EPS was extracted and evaluated for its composition. Results indicated that in the presence of 25 mM Ca^2+^ in TGY medium for the *D. radiodurans* (Kan^R^) biofilm growth had a substantial effect on the composition of the biofilm matrix. Fig. 4A shows that there was a considerable increase in the protein content, however, the increase was not statistically significant (P >0.05). Results also show that there was a slight decrease in the total eDNA (Fig. 4B), which was also found to be statistically insignificant (P >0.05). On the contrary, the production of exopolysaccharide, which was indirectly estimated in terms of total glucose concentration present in the EPS, was significantly enhanced (P< 0.05), when *D. radiodurans* (Kan^r^) biofilm was grown in the presence of Ca^2+^ as compared with the control (Fig. 4C).

**Figure 4:**
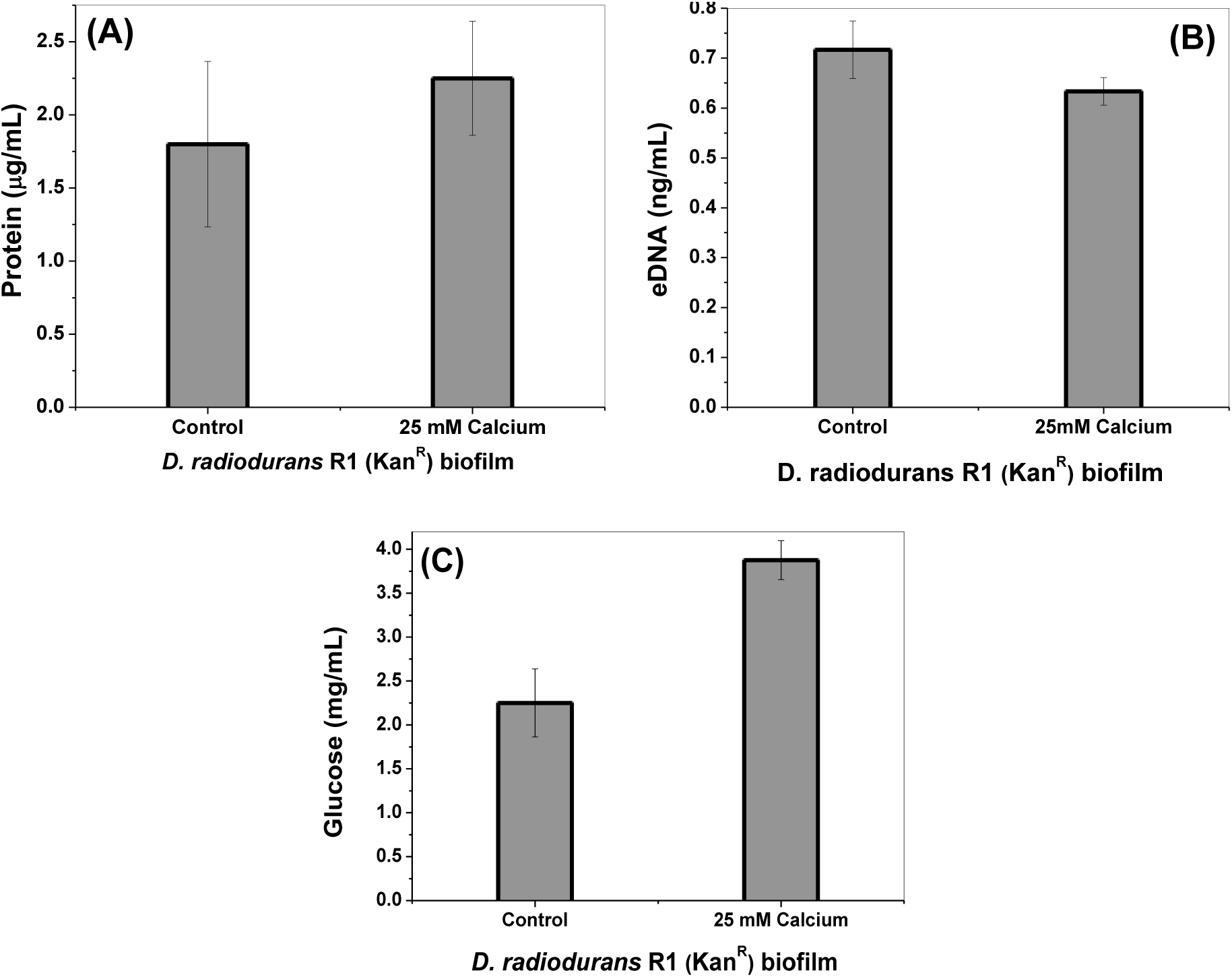
Effect of the presence of the calcium on the different constituents of extracellular polymeric substances (EPS) in *D. radiodurans* (Kan^r^) biofilm.

### Heavy metal remediation by *D. radiodurans* (Kan^r^) biofilm

*Deinococccus radiodurans* (Kan^r^) biofilm was grown for 48 h in the presence (test) and without 25 mM Ca^2+^ (control). Thereafter, the 48 h old biofilm was treated with 1 ppm aqueous solution of Co^2+^ and Ni^2+^. After 24 h of contact time, the supernatant samples were collected and Co^2+^ and Ni^2+^ concentration was analysed for the heavy metal presence using ICP-AES. Results showed that there was very less removal of Co^2+^ (~7%) and Ni^2+^ (~3%) by the *D. radiodurans* (Kan^r^) biofilm, when grown without Ca^2+^ in the growth medium (Fig. 5). Interestingly, when *D. radiodurans* (Kan^r^) biofilm was grown in the presence of Ca^2+^, it developed significant ability to remove both the heavy metals. In the presence of Ca^2+^, *D. radiodurans* (Kan^r^) biofilm could remove ~35% Co^2+^ and 25% Ni^2+^ from the 1 ppm of aqueous solution (Fig. 5). Therefore, approximately 5 fold increase was observed in Co^2+^ uptake potential, whereas ~11.3 fold increase was observed in Ni^2+^ uptake due to the presence of Ca^2+^.

**Figure 5.**
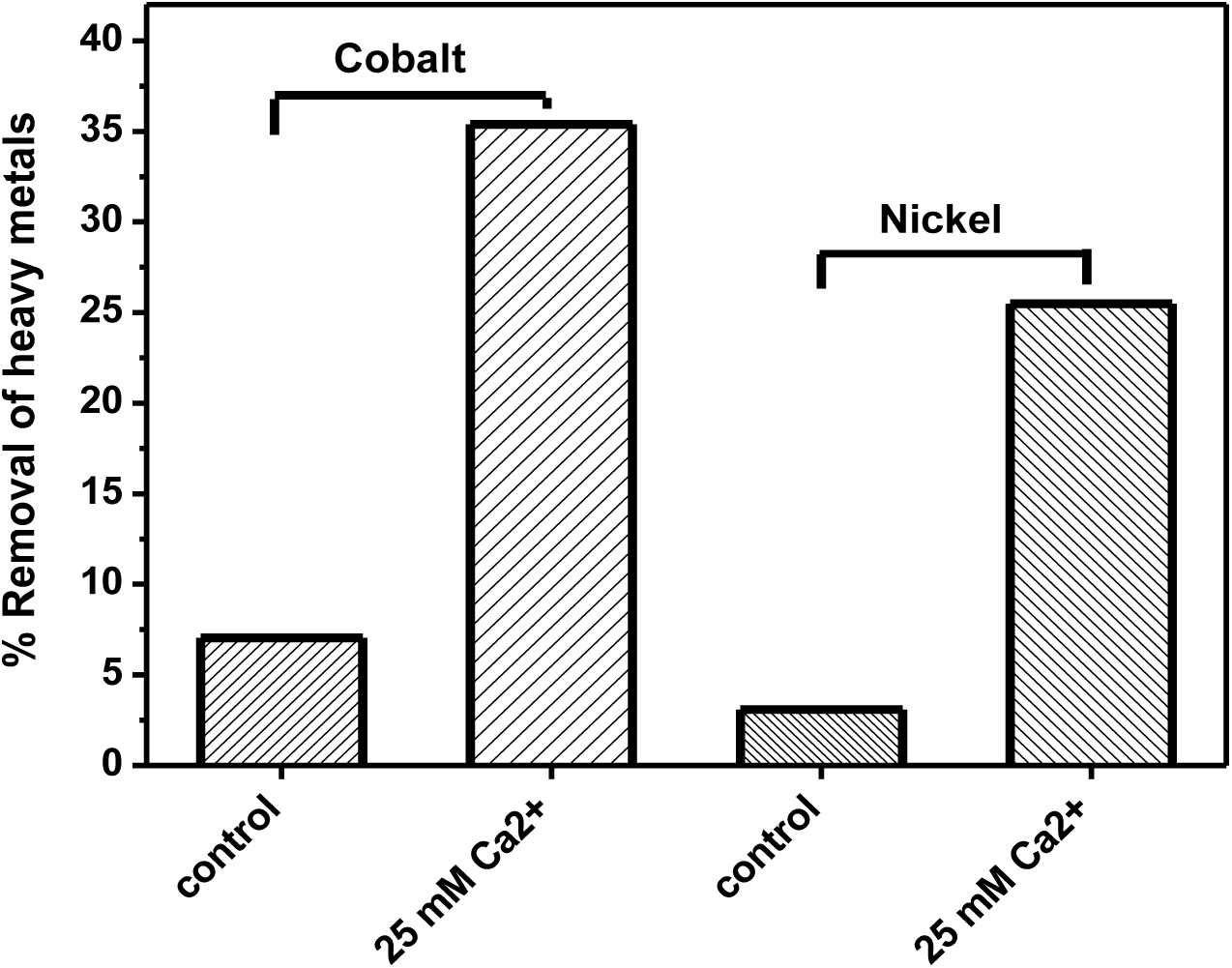
Effect of Ca^2+^ on bioremediation of heavy metals by *D. radiodurans* (Kan^r^) biofilm.

## Discussion

Nuclear industries generate radioactive heavy metal containing activated corrosion products such as ^51^Cr, ^59^Fe, ^58^Co, ^65^Zn, ^54^Mn and ^60^Co in wastes. Among the many radioactive heavy metals, ^60^Co is the most significant radioisotopes present in the activated corrosion products due to its longer half-life (5.26 Years) (Charlesworth 1971). To develop any bioremediation process to remove such radioactive element the organism needs to be highly radio-tolerant which can not only survive but can grow in chronic radiation exposure. Studies have shown that *D. radiodurans* can survive in chronic exposure of radiation, making it the most desirable bacterium to develop any bioremediation process to deal with radioactive waste. However, *D. radiodurans* R1 wild type does not form biofilm which limits its use in bioremediation despite having extreme radio-tolerance and heavy metal tolerance. *Deinococcus geothermalis,* another member of radio-tolerant *Deinococci* family, is another potential organism to develop bioremediation of radioactive heavy metals. Although few studies have shown the *D. geothermalis* forms biofilm (Kolari *et al.* 2002), but no study has indicated its potential application in bioremediation. Moreover, optimum temperature for *D. geothermalis* growth is 50°C which is not a feasible condition to maintain *in situ* applications (Makarova *et al.* 2007). On the other hand, many reports are available in the literature which shows promising potential in *D. radiodurans* as an ideal bioremediation candidate. Apart from having high heavy metal tolerance and radio-resistance, optimum growth condition for *D. radiodurans* is 30-32°C which implies that it is fit for radionuclides bioremediation applications *in situ* at ambient temperature. The only bottleneck in its large scale application is its incapability to form biofilm, which is a prerequisite for a large scale bioreactor design and applications. In this study, serendipitously, a recombinant strain of *D. radiodurans* harbouring a plasmid with a *gfp* and *kan^R^* genes found to have a significant biofilm producing property. The underlying reason behind this observation is not yet understood and is still under investigation. However, in this study, we have focused mainly on characterisation of the *D. radiodurans* biofilm and its investigating its putative role in bioremediation of heavy metals.

In general, most of the model biofilm producing organisms such as *Escherichia coli, Staphylococcus sp., Pseudomonas aeruginosa* complete a full biofilm cycle, *i.e.*, initial adhesion, irreversible adhesion, micro-colony formation, biofilm maturation followed by a dispersal phase, in 24 h under limited nutrient conditions in microtitre plate (Shukla and Rao 2014; Shukla *et al.* 2017). Doubling time for *D. radiodurans* is 1.5 h (Chen *et al.* 2009) as compared to 30-40 min for *E. coli* (Plank and Harvey 1979). Being a slow growing organism *D. radiodurans* showed the maximum biofilm at 48 h and dispersal phase was observed after 72 h. This observation also implicates that 48 h should be optimum time for developing any biofilm-mediated bioremediation process, when the process has to be performed under limited nutrient conditions or in batches.

Confocal laser scanning microscopic study of the *gfp* tagged *D. radiodurans* (Kan^r^) biofilm was done at 24 h, 48 h and 72 h to characterize biofilm parameters, such as total biomass, maximum and average thickness, average colony size, roughness coefficient, and surface to biovolume ratio. GFP fluorescence was used to image *D. radiodurans* (Kan^r^) biofilms. Various biofilm parameters were analysed from the 3D image stacks using COMSTAT software (Heydorn *et al.* 2000). Biofilm with high roughness coefficient (R*) indicates the ability of biofilm growth through micro-colonies (Shukla and Rao 2013). For *D. radiodurans* (Kan^r^) biofilm, *R*\* was found to be lowest at 48 h as compared to 24 and 72 h, indicating more homogeneous biofilm growth across the surface. Whereas, surface to biovolume ratio, which indicate that how much biofilm surface is exposed to bulk liquid, was found to be maximum at 48 h. These both parameters (low R* and high surface to volume ratio) suggest the higher availability for surface-mediated bioremediation process such as biosorption *etc*. As most of the biofilm parameter were found to be suitable as far as biofilm mediated bioremediation is concerned, 48 h old biofilm was chosen for further experiments.

In general extracellular Ca^2+^ plays an important role in maintaining the integrity of the cell wall and strengthening the biofilm matrix by cross-linking the extracellular polymeric substances and sometimes enhances the EPS production. The biofilm matrix, *i.e.*, EPS is comprised of mainly exopolysaccharide, proteins, and extracellular DNA (eDNA)(Flemming and Wingender 2010). It was hypothesized that the presence of higher concentration of Ca^2+^ ions can increase the production of EPS by *D. radiodurans* (Kan^r^). There are many reports on effect of Ca^2+^ on biofilm growth, architecture and production of extracellular matrix, especially on synthesis of exopolysaccharides (Patrauchan *et al.* 2005; Shukla and Rao 2013; Mangwani *et al.* 2014b; Shukla and Rao 2014). EPS production and composition, in turn, greatly influence the bioremediation capacity of microbial biofilms (Mangwani *et al.* 2014b; Mangwani *et al.* 2016). Earlier laboratory studies had shown that presence of Ca^2+^ enhanced the exopolysaccharide (EPS) production in biofilm in some bacteria (Mangwani et al. 2014b).

Having shown a very good biofilm forming potential, *D. radiodurans* (Kan^r^) biofilm was tested for the heavy metal removal capacity, when grown with and without the presence of Ca^2+^. As shown in the Fig. 5, fold increase was observed in Co^2+^ uptake potential, whereas ~11.3 fold increase was observed in Ni^2+^ uptake, when the biofilm was grown in the presence of Ca^2+^. As previously shown in the Fig. 4, there was a significant enhancement in the production of exopolysaccharides in the *D. radiodurans* biofilm matrix due to the presence of Ca^2+^. Therefore, it is speculated that enhanced exopolysaccharides production, which is largely negatively charged biomolecule can aid in biosorption of positively charged heavy metals such as Co^2+^ and Ni^2+^.

Biofilms are ideal for bioremediation purpose as they need not to be separated from the bulk liquid waste, thus making the whole process more economic and feasible (Shukla *et al.* 2017). Biofilm mediated bioremediation is an ideal process by a radio-tolerant microbial biofilm to ease its further downstream processing. For example, bio-precipitation of aqueous U(VI) to insoluble uranyl phosphate precipitates are not susceptible for re-oxidation in contrast to the chemically ‘reduced’ uranium mineral like uraninite which has the tendency to re-oxidise back to more soluble U(VI) (Kulkarni *et al.* 2013). Few studies using bioengineered *E. coli,* with Ni/Co transporter (NiCoT) genes showed successful removal of trace cobalt (*Co) from aqueous solutions (Raghu *et al.* 2008; Duprey *et al.* 2014). However, these recombinant *E. coli* strains did not survive beyond 20-Gy radiation exposure, limiting its use in the treatment of radioactive waste. Another study with recombinant *D. radiodurans* with cloned NiCoT genes showed higher potential of Co^2+^ removal which was ~60% removal from 8.5 nM Co^2+^ solution, *i.e.* removal of ~5.1 nM Co^2+^ (Gogada *et al.* 2015). The present study was carried out using 1 ppm (5.9 mM Co^2+^ solution) which was 1000 times concentrated and showed 35% removal of the Co^2+^ *i.e.*, 2 mM. In other words, in this study, *D. radiodurans* (Kan^R^) biofilm-mediated Co^2+^ removal method proved to be ~400 fold more efficient.

## Conclusion

This study is the first ever report on *D. radiodurans* biofilm as well as biofilm-mediated heavy-metals remediation. *D. radiodurans* is a potential candidate for the treatment of low level radioactive waste material, because of its high tolerance to radiation. The biofilm formation by a recombinant strain of *D. radiodurans* was optimized and characterized. This study also supports the fundamental role of Ca^2+^ in biofilm growth, architecture as well as biofilm-mediated heavy metal remediation. Results of this study showed that Ca^2+^ enhances the EPS production in *D. radiodurans* biofilm which modulated the biosorption capability and enhanced the removal of heavy metals (Cobalt and Nickel). Based on this study, it can be inferred that bioremediation of radioactive waste can be achieved by *D. radiodurans* biofilm.

## Acknowledgements

Authors thank Dr. Abdul Nishad P., Water and Steam Chemistry Division, BARC facilities, Kalpakkam, for providing the ICP-AES facility.

